# omniCLIP: Bayesian identification of protein-RNA interactions from CLIP-Seq data

**DOI:** 10.1101/161877

**Authors:** Philipp Drewe-Boss, Hans-Hermann Wessels, Uwe Ohler

## Abstract

High-throughput immunoprecipitation methods to analyze RNA binding protein – RNA interactions and modifications have great potential to further the understanding of post-transcriptional gene regulation. Due to the differences between individual approaches, each of a diverse number of computational methods can typically be applied to only one specific sequencing protocol. Here, we present a Bayesian model called omniCLIP that can be applied to data from all protocols to detect regulatory elements in RNAs. omniCLIP greatly simplifies the data analysis, increases the reliability of results and paves the way for integrative studies based on data from different sources.

## 1 Background

All RNA molecules are subject to post-transcriptional gene regulation (PTGR) mechanisms, including sequence-, structure -and RNA-modification-dependent modulation of splicing, cleavage and polyadenylation, editing, transport, stability, and translation. In the regulation of PTGR RNA-binding proteins (RBPs) play an important role. Many RBPs are required for constitutive processes, such as pre-mRNA splicing, cleavage, and polyadenylation. Furthermore, cell-type specific RBPs and non-coding RNAs can regulate the flow of genetic information in more directed manners, e.g. by regulating mRNA stability or translation. The complex orchestration of RBPs upon their respective targets ultimately determines appropriate protein expression.

The complexity and importance of PTGR is underscored by the large number of RNA-binding proteins (RBPs) that have been identified in recent genomics and proteomics studies ^1^ as well as the wide range of diseases that result from genetic alterations within RBPs and/or their mRNA targets ^2, 3^. Despite this large number of human RBPs, neither targets nor function for the vast majority are well understood. Uncovering the regulatory sequence elements and important RNA-RBP interactions will be critical to interpret human genetic variation in regulatory RNA regions and in the noncoding transcripts increasingly uncovered by genome-wide deep sequenc-ing ^4, 5^.

Deep sequencing technologies have enabled the development of various new protocols for mapping interaction sites between RNA-binding proteins and their RNA target sites as well as for identifying RNA-modifications on a genome-wide scale. Therefore, it is now possible to resolve interdependencies and redundancies of binding of RBPs and ribonucleoprotein particles (RNPs) to mRNA molecules and evaluate the contribution of these interactions to gene regulation in the context of cellular metabolism, organismal development or normal and disease states ^6, 7^. Ex-perimental approaches to study genome-wide RNA-RBP interactions include different variants of cross-linking and immunoprecipitation (CLIP) protocols: high-throughput sequencing of RNA isolated by crosslinking immunoprecipitation (HITS-CLIP) ^8^, photoactivatable ribonucleoside enhanced cross-linking and immunoprecipitation (PAR-CLIP) ^9^, individual nucleotide resolution cross-linking and immunoprecipitation (iCLIP) ^10^ or enhanced CLIP (eCLIP) ^11^. Similar principles have also motivated the development of protocols to study transcript modifications such as m6A-Seq ^12^ or Pseudo-seq ^13^. These protocols all have in common that they enable sequencing of RNA-fragments that were bound by a specific RBP or carry a modification, via antibodies against the native protein or tagged transgenic RBPs.

Due to particular aspects of RBP cross-linking, a crucial difference to protocols such as chromatin immunoprecipitation (ChIP)-Seq ^14^ is that the resulting fragments contain conversions, deletions or truncations at or near the cross-linked sites. These so-called diagnostic events are therefore indicative of RNA-RBP interactions or RNA modifications and thus enable nucleotide-level identification of the binding sites. For PAR-CLIP the most common diagnostic event type is a T-C conversion, for iCLIP and eCLIP it is a truncation and for HITS-CLIP deletion. It should be noted, however, that there can be also less abundant secondary diagnostic event types at the interaction sites ^15^. Similar to ChIP-Seq, the resulting data from these protocols exhibits pileups of reads (peaks) near interactions sites. The height of peaks is influenced by factors such as the strength of binding, interaction or competition with other RBPs, local biases induced by differences in cleavage and primer efficiencies. A fundamental difference to ChIP-Seq is that the coverage at interactions sites, but to a smaller degree also at non-binding sites, is strongly influenced by the wide magnitude of RNA expression levels, i.e. the relative abundance/availability of the transcript that was bound. In order to estimate the extend of confounding of the peak height by factors apart from the binding strength, input or background libraries that include most steps of the CLIP protocols except the IP can be generated. These libraries can subsequently be used to estimate other overall effect of confounding factors on the peak height. Another challenge of the data is that there are often spurious peaks at locations that do not show the typical characteristics of binding sites (e.g. motifs). These can be due for example to spurious small RNAs (e.g. miRNAs) in the sequencing libraries. In summary, the challenge of CLIP data analysis includes the proper modeling of peak height and the diagnostic events, while accounting for confounding factors and the modeling of technical and biological variance.

Various methods have been proposed to recover the interaction sites from sequencing data ^16, 17^. PARalyzer, the first dedicated tool for PAR-CLIP data analysis, mapped sites via local maxima of kernel-smoothed profiles of T-C conversion events, the most prevalent diagnostic event in PAR-CLIP data. WavCluster ^18^ models the T-C conversions and sequencing errors using a binomial distribution and estimates a background threshold to identify peak boundaries. The binomial model of T-C conversions is extended by BMIX ^10^) to also model sequence variants. Methods that do not model the diagnostic events include Piranha ^19^, which determines bins of fixed size that have a higher number of read starts than expected by chance. Piranha was the first method to model the CLIP-reads using a Negative binomial distribution and principle also allows including covariates. Another method that does not use the diagnostic events is Clipper ^19^, which models background read-counts using a Poisson distribution and identifies regions that are higher than expected by chance. However, all these methods suffer from at least one of the following shortcomings: (1) They do not contain an explicit model for diagnostic events, or they can be only applied to only a specific CLIP protocols as the modeling of diagnostic events is restricted to one of the event types. (2) They do not allow accounting for confounding factors, e.g. the gene expression. This can lead to a high false positive rate of peaks in highly expressed genes, while at the same time missing peaks for lowly expressed genes. (3) As many early datasets did not provide background or input control libraries, many tools did not allow for integration of such data. Most tools also cannot handle replicate data and thus cannot account for biological variance, leading to poorly calibrated methods.

## 2 Results

### Method overview and results

To address the shortcomings of existing methods, we developed a new Bayesian method (omniCLIP) to identify regulatory regions from all of the aforementioned protocols. The basic principle of our model is to identify target sites via an unsupervised segmentation of the transcriptome. omniCLIP learns the relevant diagnostic events directly from the data and automatically uses it during peak calling. Furthermore, it explicitly accounts for confounding factors as well as technical and biological variance (see Figure 1). To achieve this, we employ a non-homogeneous hidden Markov model (NHMM) to segment the genome into peaks and non-peaks. The emission probability of the NHMM is given by the product of the joint probability of the coverage in all replicate CLIP and background libraries and the probability of the observed diagnostic event frequency. To model coverage, we use a Negative Binomial based generalized linear model that models both confounding by the gene expression, confounding of local effects and also allows to account for excess variance. The diagnostic events are modeled using a mixture of Dirichlet-Multinomial distributions. The transition probabilities of the model are based on a logistic transfer function that depends on the coverage. All parameters of the model are learned from the data, making it easily applicable to data from various protocols.

**Figure 1:**
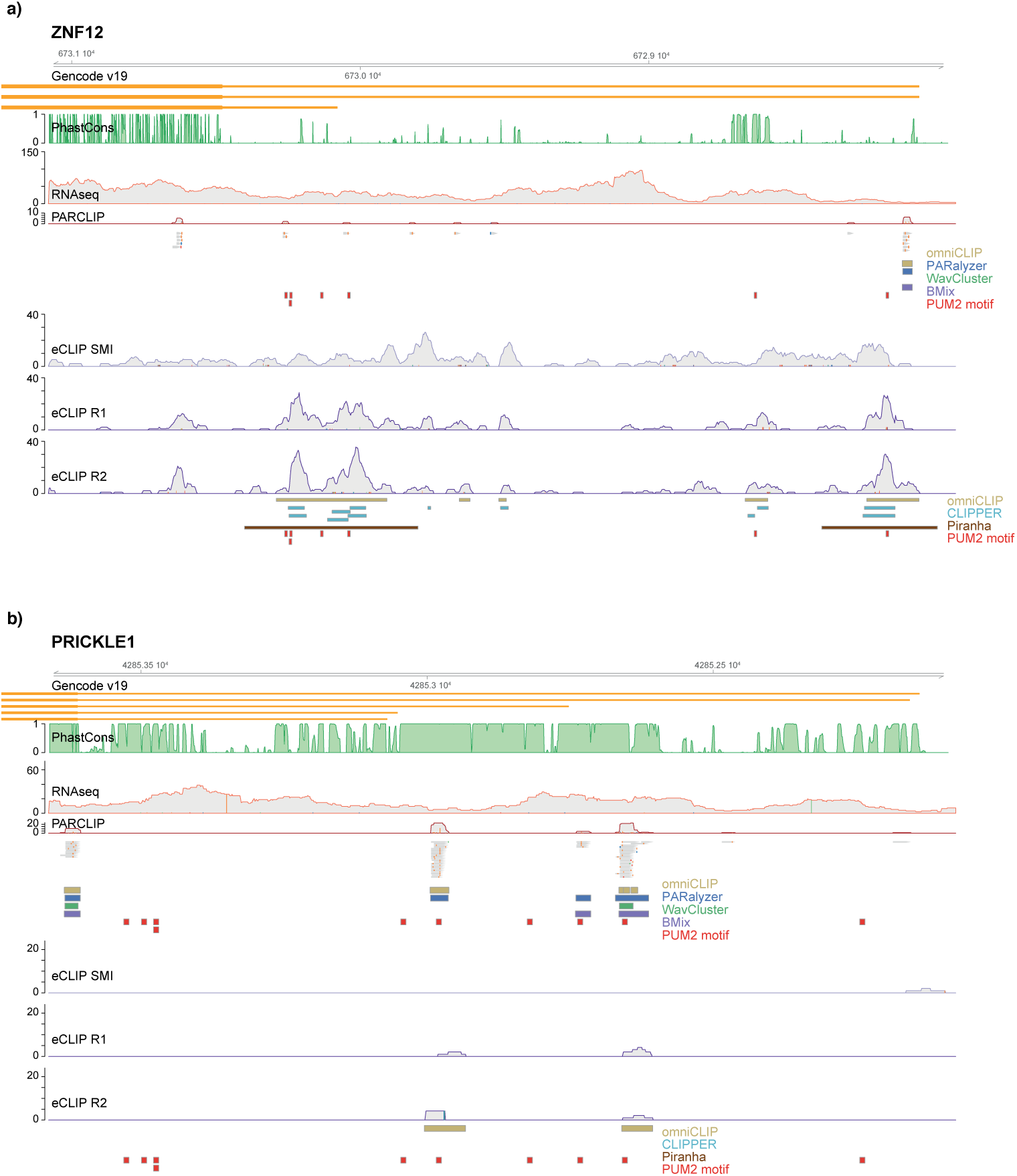
Examples of peak finding across protocols. Peak calling for the two transcripts ZNF12 and PRICKLE1 for PUM2 eCLIP libraries with a matched input from K562 cells, and for PUM2 PAR-CLIP libraries from HEK293 cells with an RNA-Seq background. On the top of each panel, Gencode v19 transcript isoforms are illustrated, as well as UCSC hg19 100way PhastCons conservation scores (green). Peak calls for omniCLIP (yellow), PARalyzer (black), WavCluster (green), BMix (purple), Clipper(turquoise) and Piranha (brown) are shown below the coverage profiles. PUM2-motifs with score higher than 8.0 are shown under the peak calls (in red). **a** The 3’UTR of ZNF12. omniCLIP can be used to robustly calls peaks on PUM2 CLIP data to determine cell type specific binding events, despite the data being generated by different specialized CLIP protocols and in across cell types. For PAR-CLIP data, individual read alignments (grey bars) are depicted to illustrate PARCLIP specific T-C conversions (organge ticks) relative to the reference genome. Due to their low depth the two PAR-CLIP libraries have been merged for the visualization. omniCLIP does not call a peak for the leftmost read cluster. This shows that the accurate modelling of diagnostic events as well as incorporation of the local background enables a better distinction of true high-confidence from low-confidence binding sites. **b** The 3’UTR of PRICKLE1. omniCLIP calls ‘true’ binding sites in regions with sparse data, highlighting the benefit of using replicate information and having a well calibrated gene expression model.

**Figure 2:**
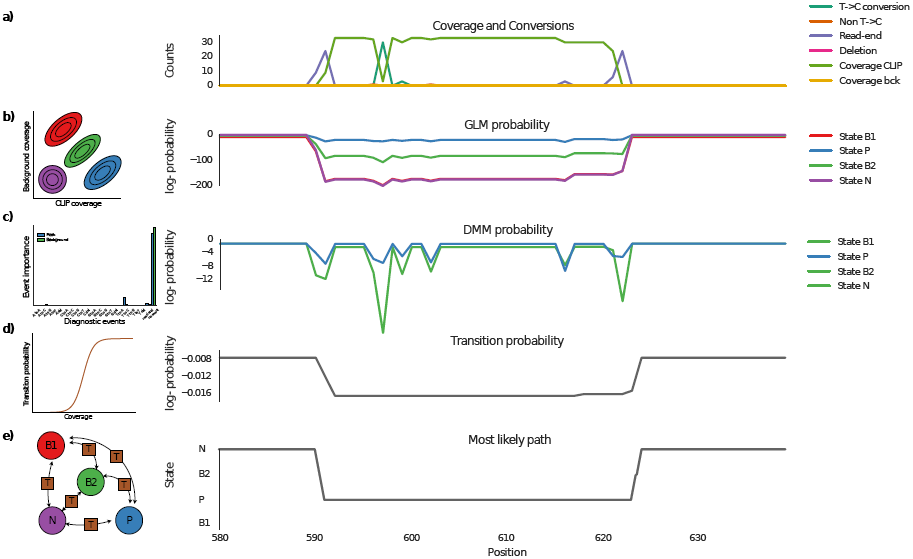
Method overview. **(a)** Coverage as well as diagnostic events. Here, the diagnostic events are subtracted from the CLIP coverage. The emission probability is computed as the product of two components. **(b)** The coverage model is based on a GLM and a model for the diagnostic events (shown in (**(c)**) that is based on a Mixtures of Dirichlet-Multinomial (DMM) distributions. **(d)** The transition probability is computed from a logistic function of the coverage. The emission and transition probabilities are used in a Non-homogeneous Hidden Markov Model to segment the sequence in to peak regions (P) and non-peak regions (N, B1, B2).

To showcase the versatile abilities of omniCLIP, we demonstrate its application across data from different CLIP protocols, for RBPs that enable an independent evaluation of the quality of peak calls. First, we assessed its performance on PAR-CLIP ^9^ and eCLIP experiments for Pumilio 2 (PUM2), a binding factor with known and high sequence specificity. To this end, we compared the predictions with those from other PAR-CLIP methods, including PARalyzer, WavCluster, Piranha, and BMIX. On this PAR-CLIP dataset obtained from the human HEK293 cell line, omniCLIP followed by PARalyzer called the highest number of peaks (*n* = 7, 900 and *n* = 5, 602, respectively) followed by BMix (*n* = 4, 501), WavCluster (*n* = 2, 473) and Piranha (*n* = 678). As there is no matching PAR-CLIP background dataset available for PUM2, we used two HEK293 ribo-zero RNA-Seq libraries as background ^7^. To evaluate the quality of the called peaks, we analyzed the enrichment of high-scoring PUM2 motifs, which we take as indicators of high-affinity binding sites. As the number of peaks called by different methods varied by an order of magnitude, we compared the enrichment only in the top 1,000 peaks of each method. For methods where no ranking criterion was provided, we used a random sub-selection of peaks. omniCLIP and PARalyzer had the highest enrichment of high-scoring PUM2 motifs in the peaks (see Figure 3(a)), and the difference to the other methods was especially strong for peaks that had a high motif score. The enrichments are not due to chance (see Supplemental Figure 1).

**Figure 3:**
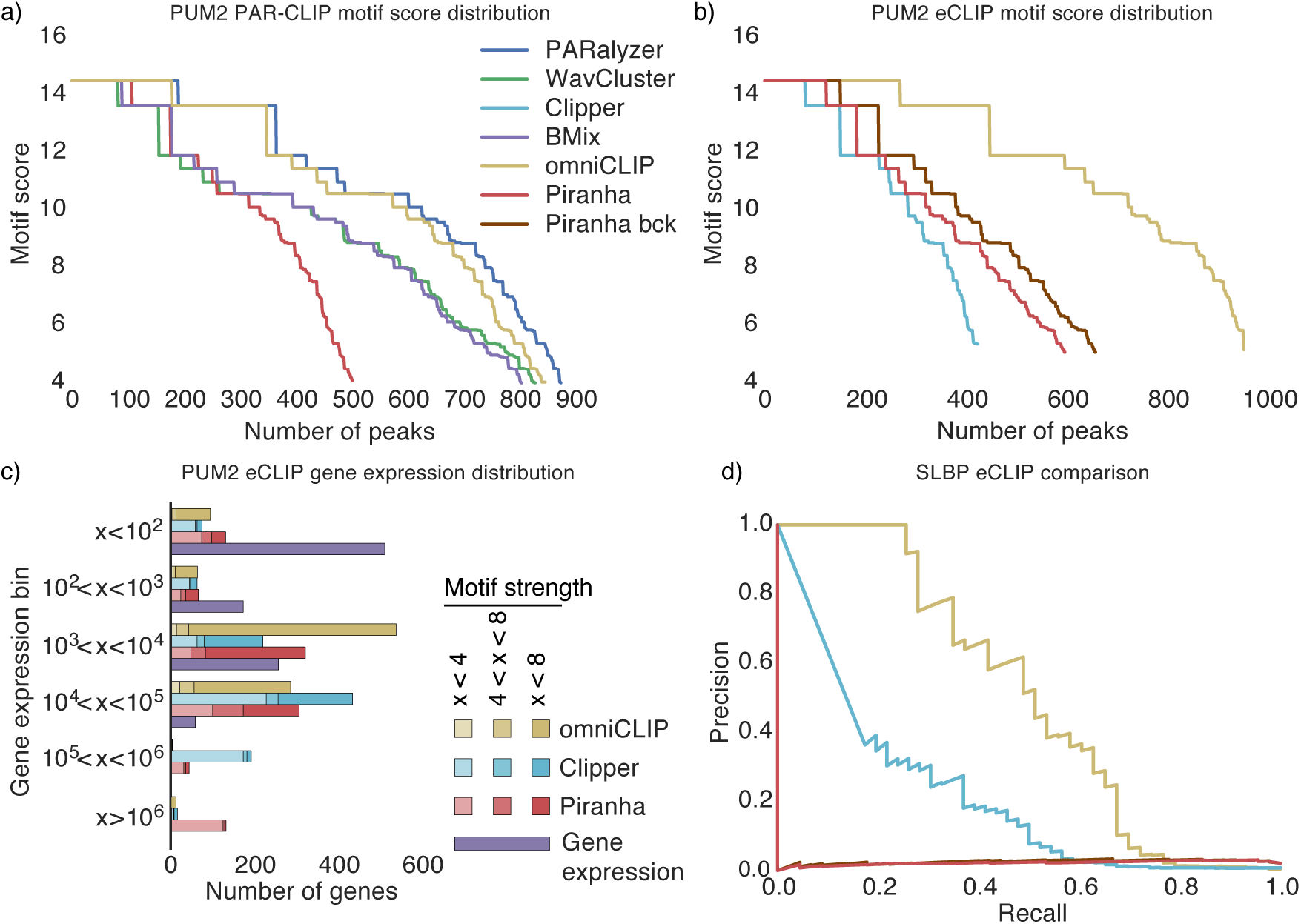
Performance evaluation. (**a**) Distribution the PUM2 motif scores of the top 1,000 called peaks for PARalyzer, Piranha, WavCluster, BMix and omniCLIP on a HEK293 PUM2 PAR-CLIP dataset. (**b**) PUM2 motif scores distribution for the top 1,000 called peaks for omniCLIP, Clipper and Piranha on a HepG2 PUM2 eCLIP dataset. (**c**) Gene expression distribution for the top 1,000 peaks on the HepG2 PUM2 eCLIP dataset as well as the expression of 1, 000 randomly sampled genes. Peaks are further classified into weak score (PUM2 PWM score *x <* 4.0), medium score (4.0 *< x <* 8.0) and high score (*x <* 8.0). On the (right) side of the plot, peaks with a PUM2-motif score of at least 4.0 are shown. On the (left), peaks without such a motif are shown. The gene expression of 1, 000 randomly sampled genes with at least 1 read count is shown for comparison in (purple). (**d**) Precision recall curves for Clipper and omniCLIP on a HepG2 SLBP dataset for discriminating histone genes and non-histone genes from peak scores.

We then applied omniCLIP to a PUM2 eCLIP dataset from the human K562 cell lines that we obtained from ENCODE. Here, we compared omniCLIP with Clipper and Piranha. We applied Piranha with and without providing it the background as a covariate. Applying Clipper results in on average 43, 594 peaks per replicates, whereas omniCLIP found 57, 634 peaks and Piranha only 17, 587 peaks, with omniCLIP exhibiting the highest enrichment of high scoring motifs in the top 1000 peaks (see Figure 3(b)). Again the enrichment of high scores in the top 1000 peaks was not due to chance (see Supplemental Figure 3). To determine how gene expression influenced the ability to detect peaks (sensitivity) as well as the quality of the detected peaks, we binned the top 1,000 peaks based on the expression level of the transcript in which they were identified (see Figure 3(c)). We further classified the peaks by whether they had a PUM2 motif with a weak score (*x <* 4.0), medium score (4.0 *< x <* 8.0), or high score (*x <* 8.0).With increasing gene expression levels, the sensitivity of peak calling increased, i.e. more peaks were found. However, the rate of peaks without strong motifs also depended on gene expression levels: For peaks in genes with less than 10, 000 counts, omniCLIP, Piranha and Clipper peaks contained 87% (860 of 983), 52% (434 of 824) and 43% (346 of 792) high scoring motifs, respectively. This was very different for peaks in genes with more than 10, 000 counts: Here, 76% (13 of 17), 6% (11 of 176) and 9% (18 of 208) of omniCLIP, Piranha and Clipper peaks had high scoring motifs. This suggests that omniCLIP has a better calibration for the peak quality than Clipper and Piranha.

Available eCLIP data for SLBP allowed for a different independent validation of peak calls, as it is known to bind specifically the 3’-ends of histone-gene mRNAs. Thus, peaks in histone transcripts should have a higher score than those found in other transcripts. Therefore, we combined the scores of all peaks in a gene and measured via the area under the precision-recall curve (auPRC), how well the scores allow distinguishing of histones from other genes. Here, omniCLIP achieved an auPRC of 0.50, Clipper an auPRC of 0.21, and Piranha an auPRC of 0.03 and 0.02 with and without using the background CLIP data, respectively (see Figure 3(d)).

To demonstrate that omniCLIP can also be used to analyze HITS-CLIP data, we applied it on libraries for the *Drosophila* RBP CNBP (CG3800), which we had previously identified as an unconventional RBP ^20^. CNBP binds mainly to mature mRNA sequences in *Drosophila* and *hu-man* ^20, 21^. Within these sequences, CNBP shows a slight preference for binding of start and stop codon proximal regions, relative to input (see Figure 4(a)) Both *Drosophila* CNBP HITS-CLIP replicates come with size matched UV-crosslinked input control of digested total RNA, collected prior to immunoprecipitation. Importantly, input RNA fragments undergo a library cloning procedure very similar to CLIP, including RNA fragment size selection and adapter ligation, resulting in highly accurate backgrounds. Application of omniCLIP resulted in 34, 224 peaks. The peaks show increasing annotation to start and stop codon categories with increasing site scores (see Figure 4(b)). This is in agreement with human CNBP, which was recently shown to bind preferentially to regions close to start codons ^21^. We identified the highly significant GGAGGA motif relative to dinucleotide shuffled background (see Supplemental Table 1) in omniCLIP peaks annotated to be mature mRNA sequences (see Figure 4(c)). This confirms the reported kmer-enrichment relative to input in concurrent *in vitro* and *in vivo* studies of the human CNBP ortholog ^21, 22^. We saw a strong connection of the motif residing in proximity to the peaks summit (see Figure 4(d)), suggesting that omniCLIP can reliably resolve biologically relevant interaction sites in HITS-CLIP data, even with lower frequencies of diagnostic events.

**Figure 4:**
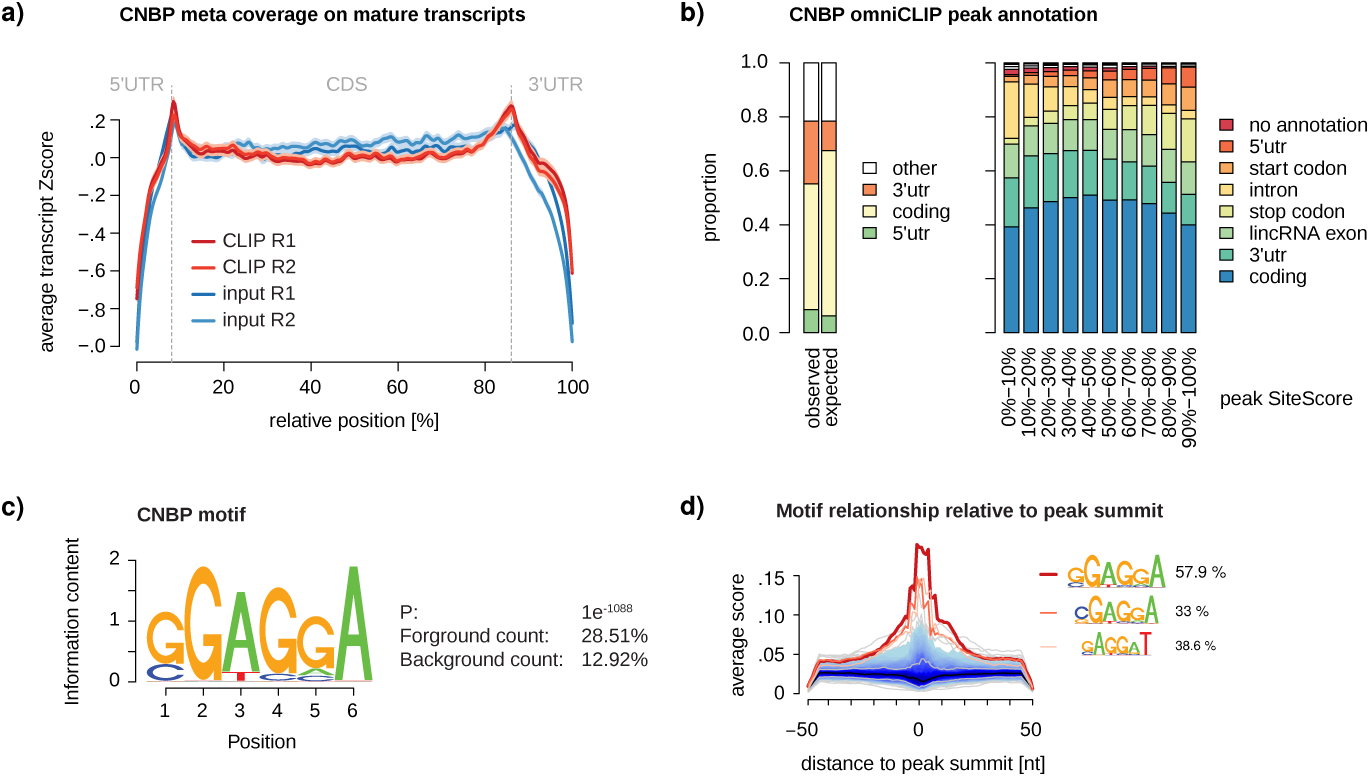
Binding preferences of CNBP. (**a**) Metaplot depicting the average Z-score transformed binned coverage across all genes (transcript with highest RSEM isoform percentage selected) with omniCLIP peak. Median 5’UTR (8%), CDS (78%) and 3’UTR (14%) proportions were extracted from all expressed genes in *Drosophila* S2 cells (TPM*>* 0) from regular total RNA-Seq experiments. Shades around solid lines indicate the standard error. (**b**) omniCLIP peak annotation grouped by strength into 10 peak SiteScore bins. (Left) Simplified annotation categories, to enable comparison to expected annotation distribution. Here, 5’UTR contains the start codon and 3’UTR the stop codon, respectively. The expected peak annotation distribution was calculated according to the feature distribution shown in (a), for all peaks tat are annotated as mature transcripts. Peaks classified as ‘other’ were ignored. (Right) Peak annotation categories grouped by peak score. Peaks annotated with start or stop codon do overlap such features. (**c**) CNBP motif calculated using HOMER2 for all peaks annotated to mature transcripts (*n* = 29556), relative to 10x dinucleotide shuffled background sequences. (**d**) Recovery of the CNBP motif and shuffled PWM relative to peak summit of all peaks used (n = 29,433). PWM match required 80% similarity. Indicated percentages reflect peak sequences with motif hit. The next highest recovered random PWMs are variants of the identified motif.

For most available datasets, we do not have meaningful, high-quality benchmark data, as many RBPs have been studied comparatively recently and motif descriptions or knowledge of the precise set of target transcripts are frequently lacking. Often, we do not even know yet which specific PTGR processes they control. To still apply omniCLIP in a more comprehensive study, we made use of ENCODE data and investigated whether splicing rates of splicing events near RBP-binding sites changed upon knock-down of the RBP ^23^. To this end, we focused on a set of splicing related RBPs for which corresponding shRNA RNA-Seq knockdown experiments were available (see Supplemental Table 2). Despite a smaller advantage of omniCLIP over Clipper (see Figure 5) compared to the above in-depth studies, the predictive value (area under the precision-recall curve) for differential splicing was consistently higher (0.25 versus 0.24). The performance difference was present at all different types of splicing events that we studied (see Supplemental Figure 2).

**Figure 5:**
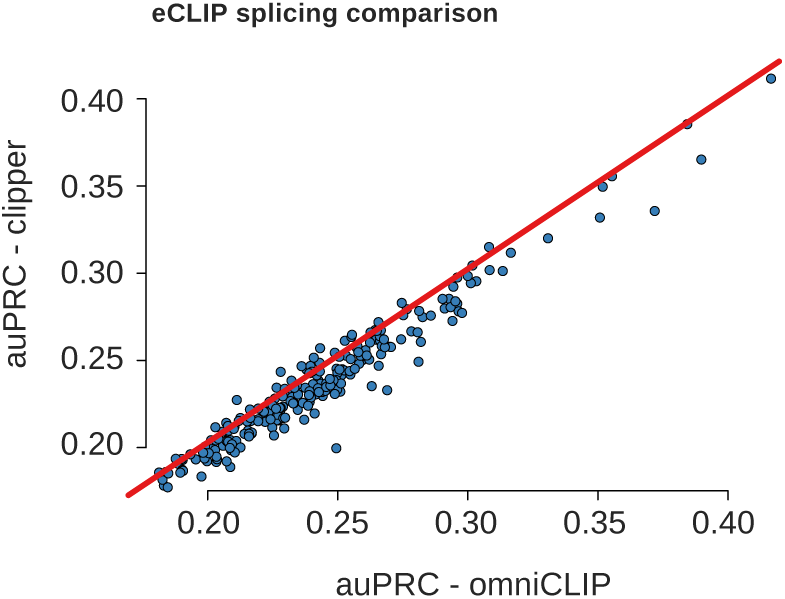
ENCODE splicing analysis. Shown is a comparison of omniCLIP and Clipper on splicing associated RBPs using eCLIP and shRNA RNA-Seq data. Datapoints are the auPRCs of omniCLIP and Clipper for predicting differential splicing events upon knock-down of a RBP based on the eCLIP peaks for the respective RBP.

## 3 Conclusions

The ability to understand the mechanisms of RNA-processing and their role in development or diseases requires understanding RBP-RNA interactions and functional consequences of these interactions. This relies on reliably identifying RBA-RNA interaction sites. However, determining the interaction sites from CLIP-Seq data is challenging due to the presence of many confounding factors.

Here, we present omniCLIP, a Bayesian approach to identify regulatory elements from CLIP data. Our model presents a principled framework for RNA interaction assays and takes into account several important new aspects. First, we quantitatively model the observed coverage in all replicates. This allows both, for including information from replicates and also accounting for various confounding factors. Second, we use an empirical Bayesian approach to identify and model important diagnostic events and sequencing errors. Finally, we take both biological and technical variance in to account in our model. We show that omniCLIP can be applied to data from a wide-range of CLIP-protocols and is straightforward to adapt to new protocols, as all parameters are learned from the data. For instance, as CLIP-Seq protocols are conceptually similar to RNA modification sequencing, omniCLIP should be easily adapted to identify RNA modifications. Finally, omniCLIP models the data in a principled way, i.e. each of its components has a clear Bayesian interpretation. This enables an easy integration of other probabilistic models in omniCLIP, such as for binding motif, structure, for various biases or explicit models of additional confounding factors.

In omniCLIP, the data used for the background modeling data plays a crucial role for measuring global and local biases. It is also used to calibrate the diagnostic event model. In general we recommend using an input as a background dataset. Yet, in many especially early published CLIP studies, these data were not acquired. In this situation, less specific data such as RNA-Seq data can serve as a substitute to some extent, but local biases cannot be modeled using this data and also the diagnostic event model may be less accurate. In the case when a specific background or input dataset is not available, we recommend to trim reads prior to alignment to match CLIP-read lengths in order to increase the similarity to CLIP-data. In general, an important factor for a reliable detection of RBP-RNA interactions is having a high quality alignment. For this, we suggest to remove multi-mapping reads and to use a stringent cut-off on the number of mismatches.

In summary, we have evaluated omniCLIP on various datasets for which either high-quality motifs are available, for which the target genes are known, or for which we have knock-down data. In all of these scenarios, we show that the omniCLIP performance is at least comparable or better than each method that we have compared it against. This is insofar remarkable as most competitor methods are tuned for specific protocols, and underlines omniCLIP’s potential for integrative transcriptome studies.

## 4 Methods

### Diagnostic event model

To represent diagnostic events and sequencing errors, we use the following model. We assume that peaks are a mixture of several classes of positions that have distinct rates of diagnostic events. In our model we have found that 10 classes are typically enough. For each of the classes we model the counts using a Multinomial-Dirichlet hierarchical model. In this model the diagnostic events, in all replicates at a given position, are assumed to be distributed according to a multinomial distribution with parameter *p*. Here *p* models the rate of diagnostic events. This parameter is at each position identical in all replicates. To allow variation in the rates between positions in the same class as well as for technical variance, we model *p* to be drawn from a Dirichlet distribution with parameter *α*. The resulting model is described in the following. Denote by *N ^j^* the number of reads covering a position *p* in replicate *i* ∈ {1, …, *I*} of the CLIP-libraries. Denote furthermore by 
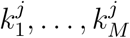
 the number of occurrences for each of the *M* diagnostic events (all possible conversions, deletions of all bases and reads ends) in the reads at position *p* in replicate *j*. If we define 
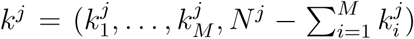
, then the probability of observing *p*(*k*^1^, *…, k^R^*) is given by:

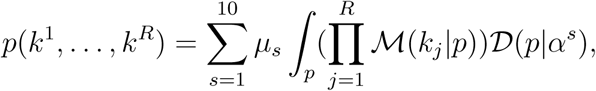

where the parameters 
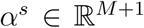
 and 
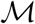
 and 
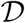
 denote the multinomial and Dirichlet distribution, respectively. The parameters are learned by maximizing the likelihood. Parameters for the peak state are fitted on the foreground dataset on the peak positions whereas parameters for the background states are fitted on the background dataset on the peak positions. Positions that are in regions where two or more genes overlap are ignored for learning the diagnostic event parameters, as diagnostic events are strand specific and overlapping genes on the opposite strand could dilute the learned signal. To speed up the fitting we estimate the parameter on a subset of 1, 000, 000 randomly sampled positions with coverage. Furthermore, to increase the stability of the fitting, we use four random initializations from a uniform distribution and the solution of the previous iteration at each iteration of the EM-algorithm.

### Coverage profile model

We jointly model the coverage in all replicates of the CLIP- and background-dataset. For our model, we assume that the coverage at each position of the genome follows a Negative Binomial (NB) distribution that is determined by the library size, the gene expression, and whether the position is a peak. We model this dependence using a generalized linear model (GLM) in the following manner. Assume that we have *I* CLIP and *J* background datasets. Then, we model the read count 
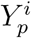
 in the CLIP-library *i* for each position *p* in a gene *g* as follows:

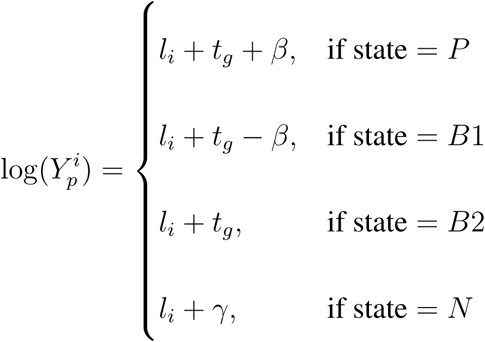

Here, *l*_*i*_ models the library size. The variable *β* models the average enrichment of CLIP-signal with respect to backgrounds in peaks, *t*_*g*_ is modeling the gene expression and *γ* models regions with little coverage (e.g. intergenic or intronic regions).

We model the coverage in the background libraries in a similar way. The read count

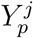

in the background-library *y* for each position *p* in a gene *g* is as follows:

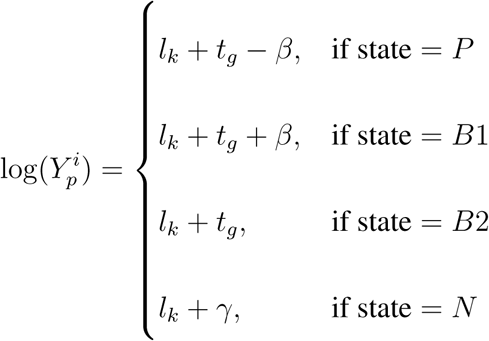

Modeling the coverage jointly across libraries allows accounting for the effect of local biases that affect the CLIP as well as the background library. For the GLM, we assume that the mean-variance relationship is described by:

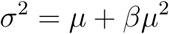

Estimation of the parameters is performed by alternately estimating the GLM-parameters *l*_*i*_, *t*_*g*_, *β* and *γ* and the over-dispersion parameter *β*. In order to ensure an equally good fit of the GLM for all states of the model, we weight the observations in each state by the inverse of the total number of observations in the state. Estimation of the GLM-parameters *l*_*i*_, *t*_*g*_, *β* and *γ* is performed using iteratively reweighted least-squares (IRLS) ^24^. In order to speed up the computation and make the solution computable in memory, we implemented an IRLS where all relevant components are sparse. For this we constrained the design matrix of the GLM such that the weighted pseudo-inverse has a sparse LU-factorization during parameter updating. This factorization in turn can be used to solve for the updated parameters. Thereby we can circumvent the computation of the pseudo-inverse, which is in general non-sparse and costly to compute.

### Modelling of the spatial dependence

To link the position-wise models for the diagnostic events and coverage profiles, we use a Non-Homogeneous Hidden-Markov model with four states. The transition-probabilities are computed based on the coverage *Y*_*p*_ in all replicates. The probability *p*_*i,j*_ of a transition from state *i* to state *j* we use:

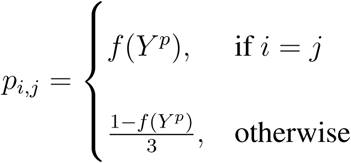

Here, we chose *f* to be the logistic function. The parameters of *f* are learned using stochastic gradient descent. To improve convergence of the GLM parameter for the background state *γ*, we set the gene expression parameter *t*_*g*_ in the computation of the emission probabilities such that all states have a higher expression rate than the background state. This is achieved by setting *t*_*g*_ = *γ* + *|β|* + 10^−5^. Adjusting these parameters is typically only necessary in the initial iterations and only for genes with few reads.

### Read filtering

To make the modeling of diagnostic events more robust, we filtered reads and masked certain positions. In order to prevent mis-mapping read ends from diluting diagnostic event profile estimation, we ignore conversions that occur in the first or last two bases of a read, and we discard reads that have more than two mismatches. Furthermore, we mask positions that are likely to be SNPs for diagnostic event modeling and calling. To this end, we use information from the background dataset to determine whether a position has a SNP. For positions to be called a SNP, we require that they have at least 20 reads and that at least 20% have a conversion event in the background.

### Peak scores

The scores for a peak are computed as the log-likelihood ratio of the peak state versus the other states in NHMM at the peak location. P-values for a peak are computed in the following way. We first compute for each position of peak the expected total coverage and variance of the CLIP-reads. For this we sum the expected mean and variance at each position of the peak. We then compute based on the CDF of a negative binomial with the computed mean and variance, the p-value of the observed total coverage of the CLIP-reads. For our analyses we only consider peaks that have Bonferroni corrected p-value *≤* 0.05.

### Model fitting

We fit the parameters of the model using the EM-algorithm. Specifically, we iterate between estimating the parameters of the diagnostic event model, the expression modeling and the NHMM. For the analyses this is done for at least 5 iterations. The model was run until full convergence was reached. As we observed that the parameters only changed minimally after 10 iterations, we stopped the model fitting after 10 iterations in order to speed up the data processing.

### Masking of miRNA genes

As a default option, we treat positions in genes that overlap annotated microRNA genes as if they had no coverage or diagnostic events.

### Data acquisition

PAR-CLIP data for PUM2 was downloaded from SRA (SRP002487). eCLIP, shRNA-Seq and RNA-Seq data for the eCLIP analysis where downloaded from the ENCODE website (https://www.encodeproject.org). HITS-CLIP data was obtained from SRA (SRP070745). Ribo-zero data for HEK293 was obtained from SRA(SRP080811).

### Read processing

Reads for PAR-CLIP analyses where processed using PARpipe (Available from https://github.com/ohlerlab/PARpipe). Reads and quantification (e.g. site calls) for ENCODE eCLIP and shRNA-Seq data were obtained from the ENCODE website (https://www.encodepro HITS-CLIP reads were quality-filtered using the fastx-tool kit using the parameters `-q 10 -p 95` ^25^ and trimmed adapters using cutadapt ^26^ using the parameters `--overlap=3 -m 24` dis-carding untrimmed reads. Subsequently, reads were transformed to fasta format and collapsed still including the 4 randomized nucleotides at both end of the reads. Randomized adapter ends got trimmed after read collapsing and added to the read identifier and treated as unique molecular identifiers (UMIs). Reads for the HITS-CLIP dataset were aligned using STAR (v.2.4.2a) ^27^. Reads were first aligned and removed against the rRNA genome parts using the following parameters for *D.melanogaster*: --alignEndsType EndToEnd --outFilterMultimapNmax 10 -outFilterIntronMotifs RemoveNoncanonical --outReadsUnmapped Fastx --alignSJoverhangMin 12 --outFilterMatchNmin 15 –outFilterMismatchNmax 1 --outFilterMismatchNoverLmax 0.05 --outFilterMultimapScoreRange 3 --alignIntronMax 20000 --seedMultimapNmax 200000 --seedPerReadNmax 30000.

The reads that did not align to the rRNA were then aligned against the *D. melanogaster* genome BDGP6 (Ensembl v81) using STAR with the following parameters: --alignEndsType EndToEnd --outFilterMultimapNmax 10 --outFilterIntronMotifs RemoveNoncanonical --alignSJoverhangMin 12 --outFilterMatchNmin 15 --outFilterMismatchNmax 1 --outFilterMismatchNoverLmax 0.05 --outFilterMultimapScoreRange 3 --alignIntronMax 20000 --seedMultimapNmax 200000 --seedPerReadNmax 30000 Reads with mismatches within the first and last two nucleotides were filtered out. Next, we removed reads with mismatches relative to the genome reference, which were likely introduced during sequencing and thus represent sequencing errors and not diagnostic events. To this end, we grouped alignments based on genomic coordinates (Chr, start, end, strand) and their UMIs. In case alignments overlapped entirely and shared the same UMI, while differing from each other and/or the reference sequence, we sorted by copy number (retained from read collapsing) and removed reads with relative lower copy number and a hamming distance one to the higher copy number reference read. For alignment of RNA-Seq reads to the *human* genome, reads were aligned against the *human* genome GRCh37 using STAR with the following parame-ters: --alignEndsType EndToEnd --chimSegmentMin 40 --chimJunctionOverhangMin 40 --outFilterMultimapNmax 2 --outFilterIntronMotifs RemoveNoncanonical --alignSJoverhangMin 16--outFilterMatchNmin 30 --outFilterMismatchNmax 2 --outFilterMultimapScoreRange 0 --alignIntronMax 20000 PAR-CLIP reads for PUM2 were aligned against the *human* genome GRCh37 using Bowtie ^28^ with the following parameters:`-v 1 -m 10 --all --best --strata -p 4 –S`

To remove reads mapping to multiple locations in our analysis, we only kept the best alignment of a read if the second best alignment had more than one mismatch more than the best alignment. Furthermore, we discarded reads that had more than two mismatches.

### Application of methods for PAR-CLIP analysis

We called peaks with PARalyzer (v1.5), Wav-Cluster (downloaded from https://github.com/FedericoComoglio/wavClusteR), Piranha(v.1.2.1) and BMIX (downloaded from https://github.com/cbg-ethz/BMix) using default parame-ters. For PAR-CLIP, peak calling with Piranha data yielded less than 10 peaks. Thus, we applied it without using a background dataset.

### Motif prediction

We predict motifs using biopython ^29^ using the pssm scoring scheme. For the motif calling a threshold score of 3.0 was used and only the forward strand was considered. Additionally a small pseudo count of 5 *∗* 10^*−*5^ was added to remove potential zeros in the PWM. The RBFOX2 position weight matrix (PWM) was obtained from ^11^ and the PUM2-PWM from ^30^.

*De novo motif discovery and visualization For de novo motiv discovery all peaks (*n* = 29556) that can be annotated by mature mRNA annotation categories (3’utr, 3’utr-intron, 5’utr, 5’utr-coding, 5’utr-intron, coding, coding-3’utr,coding-5’utr, coding-intron, intron-3’utr, intron-5’utr, intron-coding, start-codon, stop-codon) were selected. For this analysis the expressed transcripts per gene with highest RSEM isoform percentage from two total RNA-Seq experiments in *Drosophila* S2 cells (personal communications Hans-Hermann Wessels) were selected. Subsequently HOMER2 (v.4.9.1) ^31^ was used for de novo discovery using dinucleotide shuffled background sequences. For HOMER2 the following parameters were used: `len 6 -strand + -p 4`. The shuffled back-ground was generated using uShuffle (v.0.2) ^32^ using the following parameters: `-k 2 -n 10 -r 10004`. To plot the motif position relative to peak summits, we used the Bioconductor package GenomicRanges (v.1.22.4) ^33^ to center in a + -50nt window around the peak summit and searched for the motif PWM using the patternMatrix function from Genomation (v.1.2.2) ^34^ using the following parameters `min.score=0.8, prior.params = c(A=0.25, C=0.25, G=0.25, T=0.25)`. To obtain a suitable background, we randomly shuffled the PWM posterior probability from the retrieved GGAGGA motif for each nucleotide position randomly, but left the individual values unchanged to keep the overall PWM positional preference.

### Scoring for gene-based analyses

To combine peaks in for a gene we proceeded as follows. For omniCLIP we summed the scores. For Clipper and Piranha we summed the log p-values from peaks in both replicates for each gene.

### Splicing analysis

Transcript quantification for the RNA-Seq and shRNA RNA-Seq datasets of splicing related genes (see Supplemental Table2) were obtained from ENCODE. We applied SUPPA (v2.0) to determine the inclusion levels for splicing events (exon skipping, alternative 5 and 3’ splice sites, mutually exclusive exons, retained intron, alternative first and last exons) and multiple testing corrected p-values for the events ^35^. We then used a corrected p-value cutoff of 0.2 *> p* to define the true positives and 0.2 *≤ p* to define the true negatives for the computation of the auPRC.

### Software availability

The software for omniCLIP can be obtained from:

https://github.com/philippdre/omniCLIP

## 6 Author’s contributions

P.D. and U.O. conceived the project; P.D. developed the methodology with contributions by U.O. and implemented the method; P.D. and H.W. performed the analysis. P.D., H.W and U.O. wrote the paper.

## Acknowledgements

The authors would like to thank Markus Landthaler and Neelanjan Mukherjee for their input. Funding is acknowledged from DFG grant OH266/2-1 and US National Institutes of Health grant R01-GM104962.

## References

1. Gerstberger, S., Hafner, M. & Tuschl, T. A census of human RNA-binding proteins. Nature Reviews Genetics 829–845.

2. Cooper, T. A., Wan, L. & Dreyfuss, G. RNA and Disease (2009).

3. Siddiqui, N. & Borden, K. L. B. mRNA export and cancer (2012).

4. Young, R. S. & Ponting, C. P. Identification and function of long non-coding RNAs. Essays In Biochemistry 113–126.

5. Ulitsky, I. & Bartel, D. P. XLincRNAs: Genomics, evolution, and mechanisms (2013). `NIHMS150003`.

6. Mele´, M. et al. Chromatin environment, transcriptional regulation, and splicing distinguish lincRNAs and mRNAs. Genome Research 27, 27–37 (2017).

7. Mukherjee, N. et al. Integrative classification of human coding and noncoding genes through RNA metabolism profiles. Nature Structural & Molecular Biology 86–96.

8. Chi, S. W., Zang, J. B., Mele, A. & Darnell, R. B. Ago HITS-CLIP decodes miRNA-mRNA interaction maps. Nature 479–486.

9. Hafner, M. et al. Transcriptome-wide Identification of RNA-Binding Protein and MicroRNA Target Sites by PAR-CLIP. Cell 141, 129–141 (2010).

10. Ko¨nig, J. et al. iCLIP reveals the function of hnRNP particles in splicing at individual nucleotide resolution. Nature Structural & Molecular Biology 909–915.

11. Van Nostrand, E. L. et al. Robust transcriptome-wide discovery of RNA-binding protein binding sites with enhanced CLIP (eCLIP). Nature Methods 508–514.

12. Dominissini, D. et al. Topology of the human and mouse m6A RNA methylomes revealed by m6A-seq. Nature 201–206.

13. Carlile, T. M. et al. Pseudouridine profiling reveals regulated mRNA pseudouridylation in yeast and human cells. Nature 143–146.

14. Johnson, D. S., Mortazavi, A., Myers, R. M. & Wold, B. Genome-Wide Mapping of in Vivo Protein-DNA Interactions. Science 1497–1502. `20`.

15. Kishore, S. et al. A quantitative analysis of CLIP methods for identifying binding sites of RNA-binding proteins. Nature Methods 559–564.

16. Reyes-Herrera, P. H. & Ficarra, E. Computational methods for CLIP-seq data processing. Bioinformatics and Biology Insights 8, 199–207 (2014).

17. Cook, K. B., Hughes, T. R. & Morris, Q. D. High-throughput characterization of protein-RNA interactions. Briefings in Functional Genomics 14, 74–89 (2015).

18. Comoglio, F., Sievers, C. & Paro, R. Sensitive and highly resolved identification of RNA-protein interaction sites in PAR-CLIP data. BMC Bioinformatics 32.

19. Lovci, M. T. et al. Rbfox proteins regulate alternative mRNA splicing through evolutionarily conserved RNA bridges. Nature Structural & Molecular Biology 1434–1442.

20. Wessels, H. H. et al. The mRNA-bound proteome of the early fly embryo. Genome Research 26, 1000–1009 (2016).

21. Benhalevy, D. et al. The Human CCHC-type Zinc Finger Nucleic Acid-Binding Protein Binds G-Rich Elements in Target mRNA Coding Sequences and Promotes Translation. Cell Reports 18, 2979–2990 (2017).

22. Ray, D. et al. RNAcompete methodology and application to determine sequence preferences of unconventional RNA-binding proteins. Methods 118–119, 3–15 (2017).

23. Sundararaman, B. et al. Resources for the Comprehensive Discovery of Functional RNA Elements. Molecular Cell 61, 903–913 (2016).

24. Holland, P. W. & Welsch, R. E. Robust regression using iteratively reweighted least-squares. Communications in Statistics - Theory and Methods 813–827.

25. HannonLab. FASTX toolkit (2014).

26. Martin, M. Cutadapt removes adapter sequences from high-throughput sequencing reads. EMBnet.journal 10. `ISSN 2226-6089`.

27. Dobin, A. et al. STAR: Ultrafast universal RNA-seq aligner. Bioinformatics 29, 15–21 (2013).

28. Langmead, B., Trapnell, C., Pop, M. & Salzberg, S. Ultrafast and memory-efficient alignment of short DNA sequences to the human genome. Genome Biol. 10, R25 (2009).

29. Cock, P. J. et al. Biopython (2009).

30. Kassuhn, W., Ohler, U. & Drewe, P. Cseq-Simulator: A Data Simulator for Clip-Seq Experiments. Pac Symp Biocomput 433–444.

31. Heinz, S. et al. Simple Combinations of Lineage-Determining Transcription Factors Prime cis-Regulatory Elements Required for Macrophage and B Cell Identities. Molecular Cell 38, 576–589 (2010). `0801.2587`.

32. Jiang, M., Anderson, J., Gillespie, J. & Mayne, M. uShuffle: A useful tool for shuffling biological sequences while preserving the k-let counts. BMC Bioinformatics 192.

33. Aboyoun P, Pages H, L. M. GenomicRanges: Representation and manipulation of genomic intervals. R package version 1, 1–5 (2010). `arXiv:1011.1669v3`.

34. Akalin, A., Franke, V., Vlahoviček, K., Mason, C. E. & Schübeler, D. Genomation: A toolkit to summarize, annotate and visualize genomic intervals. Bioinformatics 31, 1127–1129 (2015).

35. Alamancos, G. P., Pages, A., Trincado, J. L., Bellora, N. & Eyras, E. Leveraging transcript quantification for fast computation of alternative splicing profiles. RNA (New York, N.Y.) 21, 1521–1531 (2015).

